# Temporal signal-to-noise changes in combined multiband- and slice-accelerated echo-planar imaging with a 20- and 64-channel coil

**DOI:** 10.1101/641902

**Authors:** Philipp Seidel, Seth M. Levine, Marlene Tahedl, Jens V. Schwarzbach

## Abstract

Echo planar imaging (EPI) is the most common method of functional magnetic resonance imaging for acquiring the blood oxygenation level-dependent (BOLD) contrast. One of the primary benefits of using EPI is that an entire volume of the brain can be acquired on the order of two seconds. However, this speed benefit comes with a cost. Because imaging protocols are limited by hardware (e.g., fast gradient switching), researchers are forced to compromise between spatial resolution, temporal resolution, or whole-brain coverage. Earlier attempts to circumvent this problem included developing protocols in which slices of a volume were acquired faster (i.e., slice (S) acceleration), while more recent protocols allow for multiple slices to be acquired simultaneously (i.e., multiband (MB) acceleration). However, applying such acceleration methods can lead to a reduction in the temporal signal-to-noise ratio (tSNR), which is a critical measure of the stability of the signal over time. Here we show, in five healthy subjects, using a 20- and 64-channel receiver coil, that enabling S-acceleration consistently yielded, as expected, a substantial decrease in tSNR, regardless of the receiver coil employed, whereas tSNR decrease resulting from MB acceleration was less pronounced. Specifically, with the 20-channel coil, tSNR of upto 4-fold MB-acceleration is comparable to that of no acceleration, while up to 6-fold MB-acceleration with the 64-channel coil yields comparable tSNR to that of no acceleration. Moreover, observed tSNR losses tended to be localized to temporal, insular, and medial brain regions and were more noticeable in the 20-than in the 64-channel coil. Conversely, with the 64-channel coil, the tSNR in lateral frontoparietal regions remained relatively stable with increasing MB factors. Such methodological explorations can inform researchers and clinicians as to how they can optimize imaging protocols depending on the available hardware and the brain regions they want to investigate.

## Introduction

With the introduction of echo planar imaging (EPI) for magnetic resonance imaging (MRI) by Mansfield in 1977, acquiring an entire brain volume in a matter of seconds became a reality [1]. Later, the discovery that EPI is especially sensitive to changes in blood oxygenation [2] led to EPI becoming the standard MRI protocol for investigating physiological changes in the brain both in response to stimuli [3, 4] and at rest [5].

One of the primary benefits of MRI in neuroscience research is the high spatial resolution attainable when investigating the whole brain, which, however, interacts with lower temporal resolution, whole-brain coverage, and signal-to-noise ratio (SNR). The interplay between these four properties leads to the fact that sampling a volume with an increased temporal resolution results in fewer maximum attainable slices per volume (i.e., reduced whole-brain coverage) and less acquisition time per voxel (leading to reduced signal). Conversely, sampling a volume with an increased spatial resolution results in more voxels per volume, which yields reduced signal from the smaller voxels, and a longer acquisition time may become necessary to cover the whole brain.

Multiple advances in image acquisition protocols and image reconstruction algorithms have worked toward providing both increased spatial and temporal resolution. Some of these approaches, such as partial Fourier acquisition [6] and parallel imaging, e.g. the GeneRalized Autocalibrating Partial Parallel Acquisition (GRAPPA) algorithm [7] (see also SENSE [8] and SMASH [9]), reduce the acquisition time of single slices (i.e., in-plane acceleration) and are known as slice-acceleration (S) methods. Despite their speed benefits, these approaches suffer from increased sensitivity to motion in higher acceleration factors [10–12] and the fact that SNR is inversely proportional to the square root of the acceleration factor [13].

An alternate approach to parallel imaging, called multiband (MB) imaging, consists of acquiring multiple slices at the same time by simultaneously exciting a set of slices [14–16]. Recently, a multitude of studies has investigated various aspects of MB protocols, such as the achievable acceleration factors of MB with and without in-plane acceleration [17], the effects on the temporal signal-to-noise ratio (tSNR: i.e., the mean of a voxel’s BOLD signal over time divided by its standard deviation over time) at the whole-brain level [12] and in a region-specific manner [18], the effects on g-factor noise amplification [18, 19] and signal leakage across simultaneously acquired slices [20], and their benefits with respect to the sensitivity of signal detection and statistical analysis in 3 and 7 Tesla MR scanners [21–23].

At present, there have been no systematic investigations into 1) the question of where in the brain MB- and S-acceleration (and their combinations) most prominently affect tSNR when other scanning parameters are held constant and 2) whether the type of receiver coil interacts with acceleration methods with respect to tSNR changes, as well. These outstanding concerns are critical to cognitive neuroscientists and clinical researchers who may have region-specific hypotheses to investigate using otherwise standard hardware setups. To this end, we examined these issues by scanning a gel phantom and five healthy participants with T2*-weighted EPI sequences that employed constant parameters differing only in their combinations of MB- and S-acceleration, using both a 20- and 64-channel receiver head coil. This protocol allowed us to compare the accelerated sequences to a reference scan (i.e., a scan without any MB- or S-acceleration) providing insight into the performance of different acceleration factors and where in the brain MB- and S-acceleration lead to tSNR decreases. Such results can inform researchers regarding the feasibility of different combinations of MB- and S-acceleration depending on the experimental demands and the brain regions to be investigated.

## Materials and Methods

### Participants

We acquired data for a gel phantom as well as for five human participants (1 male, 4 females; age range: 22-25 yrs, mean age [± SD]: 23.6 ± 1.52). After a standard screening to exclude contraindications for MRI, all participants provided written informed consent. All procedures were approved by the local ethics committee.

### Data acquisition

All imaging data were acquired with a 3T Siemens Prisma (Siemens Healthcare, Erlangen, Germany) using both a 20- and a 64-channel receiver head-coil (Siemens Healthcare, Erlangen) at the University of Regensburg. We used multiband (MB) sequences provided by the Center for Magnetic Resonance Research (CMRR, Minnesota) [16, 24] for all acquisitions; for slice acceleration (S), we employed Siemens’ GRAPPA [7].

All functional sequences consisted of 100 volumes covering the whole brain with 60 slices (or 64 for MB8) and 2 × 2 × 2 mm^3^ voxels employing the following parameters: time to repeat (TR) = 4800 ms; echo time (TE) = 30 ms; flip angle (FA) = 55° [20]; excitation pulse duration = 9 ms; echo spacing = 0.58 ms; bandwidth = 2368 Hz/pixel; acquisition matrix (AM) = 96 × 96; Field of View (FoV) = 192 × 192 mm^2^; partial Fourier = 6/8. The fMRI sequences differed in their MB-acceleration (MB = 1, 2, 4, 6, 8[, 10, 12 for the phantom]) and their S-acceleration (S = 1, 2; where the number indicates the GRAPPA acceleration factor). The T1-weighted Magnetization Prepared RApid Gradient Echo (MPRAGE) structural scan (not acquired for the phantom) was used for co-registration and surface reconstruction (TR = 1910 ms; TE = 3.67 ms; FA = 9°; FoV = 250 mm^2^; AM = 256 × 256). See tables 1 and 2 for a succinct description of sequence specifications.

**Table 1.** MRI acquisition parameters (*values in parentheses apply only to phantom acquisitions)

| Parameters | Sequences |  |
| --- | --- | --- |
|  | T2* EPI | T1 |
| Time to repeat (ms) | 4800 | 1910 |
| Time to echo (ms) | 30 | 3.67 |
| Flip angle ($^\circ$) | 55 | 9 |
| Excitation pulse duration (ms) | 9 | - |
| Echo spacing (ms) | 0.58 | - |
| Bandwidth (Hz/pixel) | 2368 | 180 |
| Acquisition matrix | $96 \times 96$ | $256 \times 256$ |
| Field of view (mm) | $192 \times 192$ | $250 \times 250$ |
| Voxel resolution (mm) | $2 \times 2 \times 2$ | $0.9766 \times 0.9766 \times 1$ |
| Partial Fourier | 6/8 | - |
| Multiband acceleration factors | 1,2,4,6,8,(10,12)* | - |
| Slice acceleration factors | 1,2 | 2 |

**Table 2.** Realized protocols (H: human participants, P: phantom) for both coil types (20- and 64-channel)

|  | MB1 | MB2 | MB4 | MB6 | MB8 | MB10 | MB12 |
| --- | --- | --- | --- | --- | --- | --- | --- |
| S1 | H,P | H,P | H,P | H,P | H,P | P | P |
| S2 | H,P | H,P | H,P | H,P | H,P | - | - |

For both human participants and the phantom, we separated data acquisition into two days to reduce the likelihood of changes in tSNR due to gradient heating and to maintain a similar protocol between human and non-human participants (as a single session lasted approximately 1 hour 45 minutes). For two of the five participants, we used the 20-channel coil for the first session and the 64-channel coil for the second session (and vice versa for the remaining three participants, to counterbalance the order).

For the phantom, within a given session we acquired twelve scans. The first seven scans of a session differed in their MB acceleration factors and were followed by five more EPI sequences that differed in both their MB acceleration factors and slice-acceleration (i.e., GRAPPA).

For human participants, a given session consisted of eleven scans, ten of which were T2*-weighted EPI sequences for which we instructed our participants to close their eyes, let their mind wander, and try not to fall asleep. The first five EPI scans of a session differed in their MB acceleration factors and were followed by the T1-weighted structural scan. Following the structural scan, we acquired the remaining five EPI scans in which we combined MB- and S-acceleration.

### Data analysis

Analysis of the MRI data were performed using the FMRIB Software Library (FSL) [25] version 5.0.9, Freesurfer [26, 27], MATLAB R2016a (The Mathworks, Natick, MA, USA), and CoSMoMVPA [28].

#### Phantom

Following the quality assurance protocols laid out by Freedman and Glover [29], we selected a central slice of 21 × 21 voxels from the functional data acquired from the phantom and extracted the 441 BOLD time courses (for each of the 24 combinations of MB-acceleration, S-acceleration, and coil type), from which the tSNR was computed per time course and then averaged, yielding one value per experimental condition.

#### Human participants

##### Co-registration and surface reconstruction

Functional MRI data were first preprocessed using FSL’s BET [30] for brain extraction and FLIRT [31, 32] for co-registration of each participant’s functional MRI data to their respective structural scan with 6 degrees of freedom. No spatial smoothing was performed [33]. For group-level statistics, we additionally co-registered each participants’ data to the Montreal Neurological Institute (MNI) 2mm standard space and used Freesurfer for surface reconstruction and to visualize statistical maps on inflated surface brains.

##### Statistical analysis

In order to evaluate differences in tSNR as a function of different acceleration methods, we subtracted the tSNR maps of our reference scans (i.e., those without any acceleration) from all other scans (per coil); resulting values below zero thus indicate a decrease in tSNR in the accelerated sequence (compared to the non-accelerated sequence), whereas values above zero indicate an increase in tSNR in the accelerated sequence (compared to the non-accelerated sequence). Doing so for every voxel results in a whole brain ΔtSNR map. After computing such ΔtSNR maps for each participant, we assessed group-level tSNR changes using both a whole-brain average approach and a voxelwise approach. This two-step approach allowed us first to compare a global effect to the pattern of changes observed from the phantom analysis and secondly to further localize specific changes to particular regions of the brain.

Group-level changes in global tSNR were specifically assessed by averaging all the ΔtSNR values across voxels per participant. We then submitted the resulting average ΔtSNR values to a three-way repeated-measures Analysis of Variance (ANOVA) with factors Coil (20 channels, 64 channels), MB (1, 2, 4, 6, 8), and S (1, 2).

Group-level changes in voxelwise tSNR were assessed with one-sample t-tests against zero. Due to the inflated family-wise error rate (FWER) resulting from the multiple testing procedure, we carried out an exhaustive permutation test, in which we flipped the sign of the ΔtSNR values within a given participants’ map, recomputed the t-test, and stored the resulting minimum t-score of the map [34], as we were specifically seeking decreases in tSNR. As our sample consisted of five participants, there were 2^5^ – 1 unique permutations to perform, which provided us with a null distribution of 31 random minimum t-scores per condition, to which we compared our observed t-scores, yielding FWER-corrected empirical p-values at each voxel (per condition). Furthermore, because awareness of tSNR decreases is a methodological issue, which can greatly impact the quality of one’s measurements, we feel that it is in a researcher’s interest to be statistically liberal in rejecting null hypotheses reflecting no decrease in tSNR. For this reason, we present our voxelwise analyses in two ways to increase the transparency of the underlying effects: 1) the number of voxels (per condition) that contain ΔtSNR values at four different FWER-corrected thresholds (i.e., p < 0.03, 0.1, 0.19, and 0.48, which approximately correspond to 1/31, 3/31, 6/31, and 15/31, respectively) and 2) heatmaps of uncorrected t-scores (from t_4_ = ±3, p < 0.02) projected onto cortical surfaces to depict where such tSNR decreases tend to occur.

## Results

### Most forms of acceleration decrease tSNR in the phantom

Investigating changes in tSNR as a function of different combinations of acceleration methods within a central ROI of the phantom revealed that S-acceleration led to more immediate drastic reductions in tSNR than MB-acceleration. Additionally, the 64-channel coil appears to slightly outperform the 20-channel coil, especially in the absence of S-acceleration. While lower-levels of MB-acceleration do not appear to yield extreme changes in tSNR, beyond a total acceleration of 10, all combinations of factors (i.e., S, MB, and coil type) seem to perform equally poorly (Fig. 1). These results provided a qualitative baseline for comparison when carrying out the same analysis in fMRI data acquired from the human participants.

**Fig. 1.**
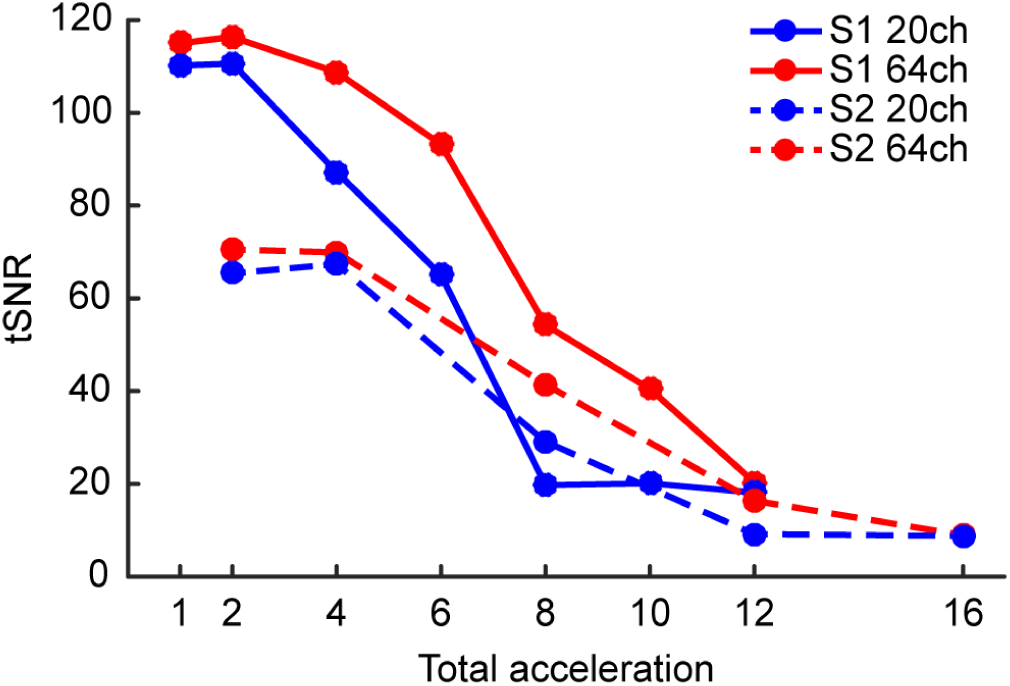
Changes in tSNR as a function of combined acceleration methods in a central region of interest (ROI) of the phantom. The x-axis represents the total acceleration factor (i.e., the product of MB- and S-acceleration). Employing slice-acceleration leads to an immediate decrease in tSNR, whereas MB-acceleration tends to yield decreases in tSNR only with higher acceleration factors. Moreover, the 64-channel coil appears to yield less tSNR decreases compared to the 20-channel coil.

### Most forms of acceleration decrease tSNR in the human brain independent of the coil

Obtaining whole-brain average tSNR values for each condition across participants yielded qualitatively similar results to those of the phantom analysis when comparing tSNR changes based on total acceleration (Fig. 2A), with the notable exception of the downward baseline shift from the tSNR values in the phantom. However, as total acceleration is generally not equivalent across acceleration methods [19], we also analyzed the tSNR results condition-wise (Fig. 2B) with a three-way repeated-measures ANOVA (Table 3). This analysis ultimately revealed that a general decrease in tSNR results from both S-acceleration (F_1,4_ = 42.4, p < 2.87 × 10^−3^) and MB-acceleration (F_4,16_ = 89.2, p < 10^−6^), though not in identical manners when the two acceleration methods are combined (S × MB: F_4,16_ = 6.35, p < 2.94 × 10^−3^). Additionally, although we were unable to find an overall difference between the two coils in terms of whole-brain average tSNR changes (F_1,4_ = 0.169, p < 0.70) and with respect to different levels of S-acceleration (Coil × S: F_1,4_ = 0.602, p < 0.48), the coils did appear to yield tSNR differences depending on the MB-acceleration (Coil × MB: F_4,16_ = 5.02, p < 8.13 × 10^−3^), in that the 20-channel coil seemed to perform slightly better with lower levels of MB-acceleration, whereas the 64-channel coil seemed to perform slightly better at higher levels of MB-acceleration. This specific pattern was marginally pronounced when additionally taking S-acceleration into consideration (Coil × S × MB: F_4,16_ = 2.26 p < 0.108).

**Table 3.** Results of the three-way repeated-measures ANOVA. Asterisks indicate effects that surpassed a statistical threshold of p < 0.05.

|  |  | F | df | p |
| --- | --- | --- | --- | --- |
| Main effects | Coil | 0.169 | 1,4 | 0.702183 |
|  | S | 42.4 | 1,4 | *0.002869 |
|  | MB | 89.2 | 4,16 | *0.000001 |
| Interaction effects | Coil × S | 0.602 | 1,4 | 0.481810 |
| | Coil × MB | 5.02 | 4,16 | *0.008127 |
| | S × MB | 6.35 | 4,16 | *0.002938 |
| | COIL × S × MB | 2.26 | 4,16 | 0.107901 |

**Fig. 2.**
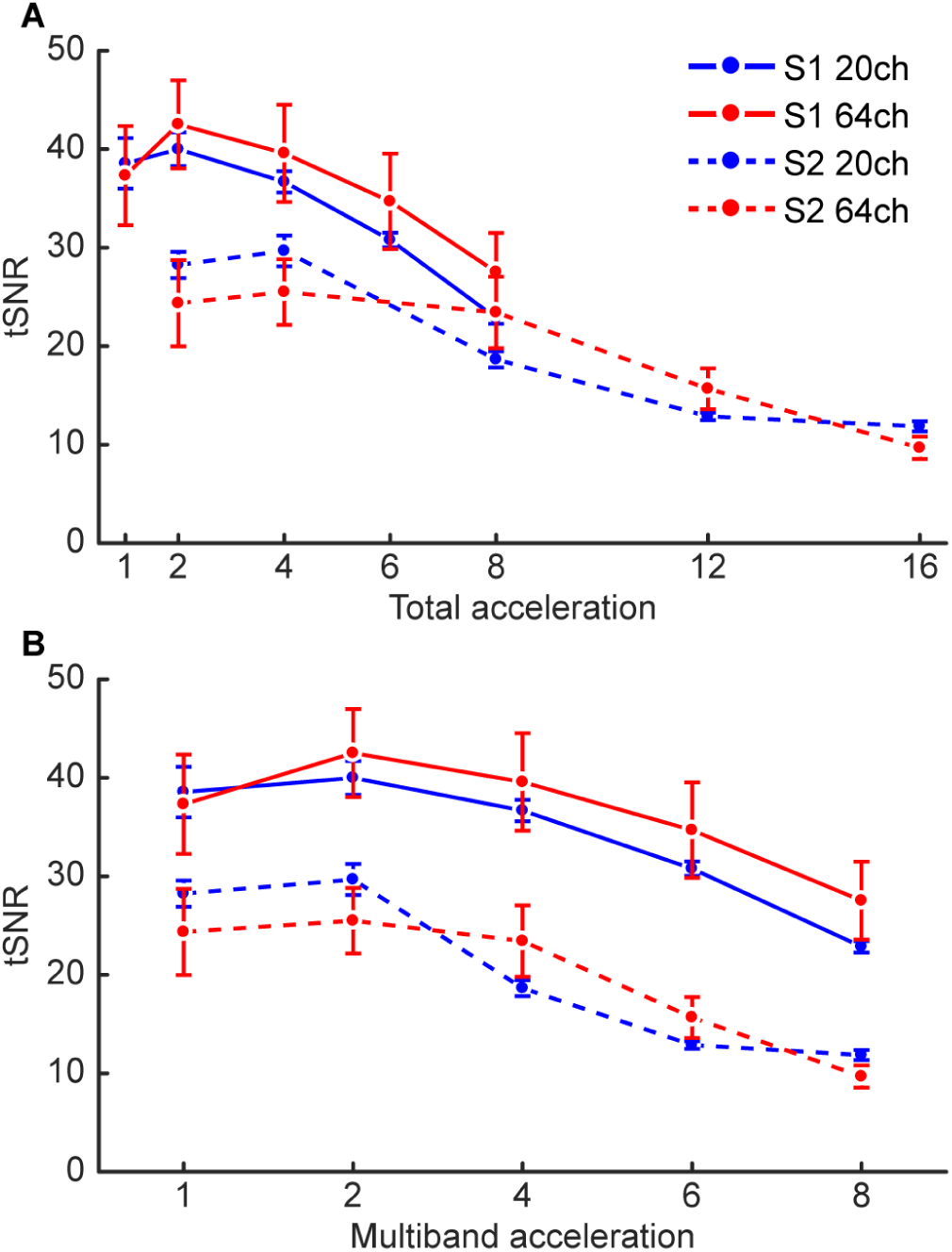
Group-level changes in tSNR from the whole-brain average analysis. **(A)** Results plotted as a function of total acceleration, as in figure 1. **(B)** Results plotted as a function of the experimental of conditions (i.e., not multiplying the MB- and S-acceleration factors). Overall, the results reflect those from the phantom analysis, namely that enabling S-acceleration leads to more substantial decreases in tSNR; however, the difference between the two coils is not as pronounced in this analysis. Error bars represent one standard error of the mean.

### Up to six-fold MB-acceleration with the 64-channel coil sustains minimal losses

While the whole-brain average tSNR provides a simple global measure, it entirely sacrifices spatial specificity of tSNR loss, which may be of critical interest depending on the experimental or clinical investigation. For this reason, we also carried out voxelwise tests of tSNR changes by comparing each combination of S-acceleration and MB-acceleration against a baseline of no acceleration (per coil). In order to balance the need for statistical rigor with the importance of detecting tSNR decreases, we present both FWER-corrected results (Fig. 3) and uncorrected results (Fig. 4, see also next section) from this analysis.

**Fig. 3.**
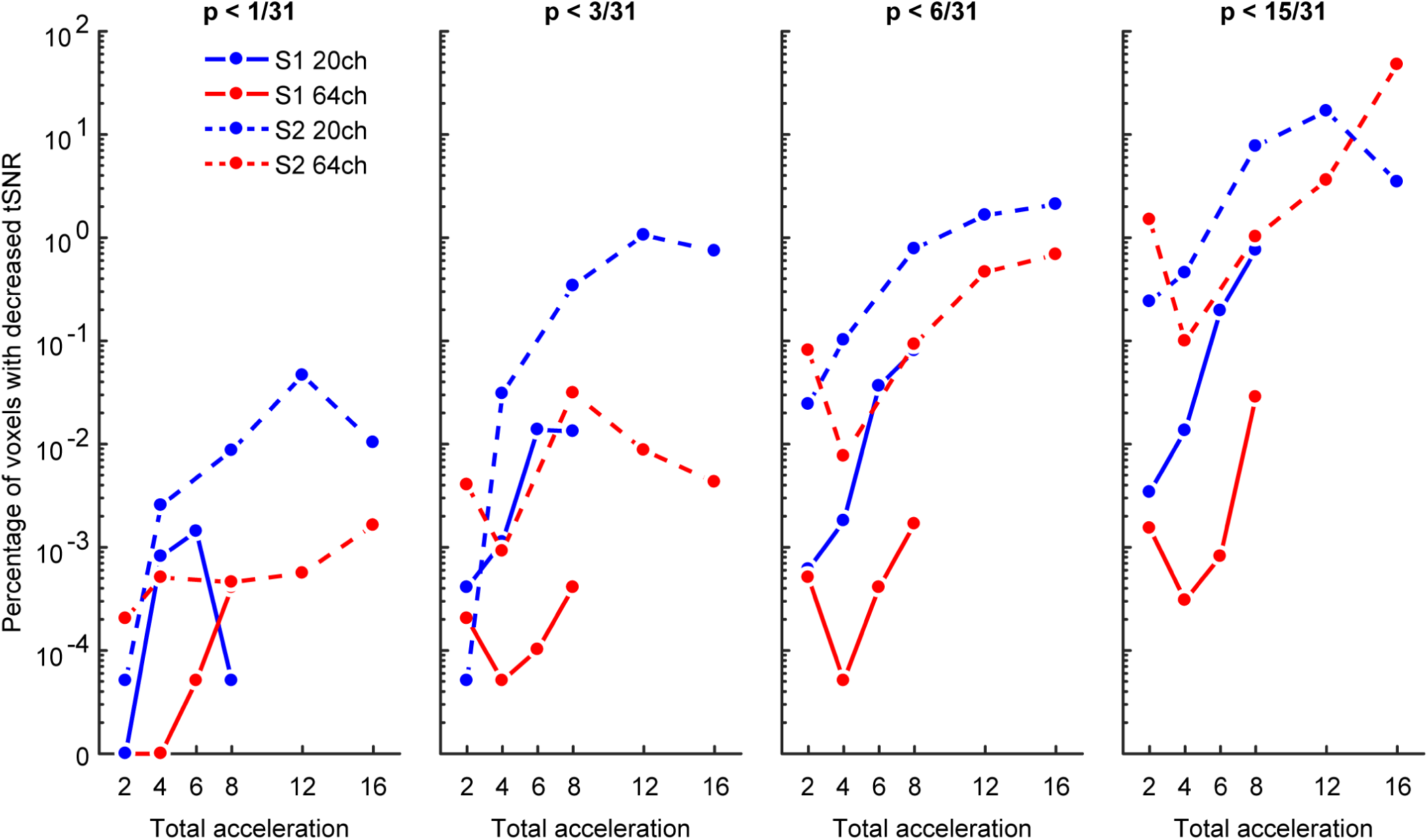
Temporal SNR decreases at each of four thresholds (f.l.t.r. p < ∼0.05, ∼0.1, ∼0.2, and ∼0.5) plotted (on a pseudo-log scale) as the percentage of voxels whose t-scores from each ΔtSNR map surpassed the FWER-corrected threshold for detecting tSNR decreases. Note that even at the more liberal FWER-corrected thresholds, the channel coil with no S-acceleration yielded the least number of voxels with tSNR decreases, especially at intermediate MB factors.

**Fig. 4.**
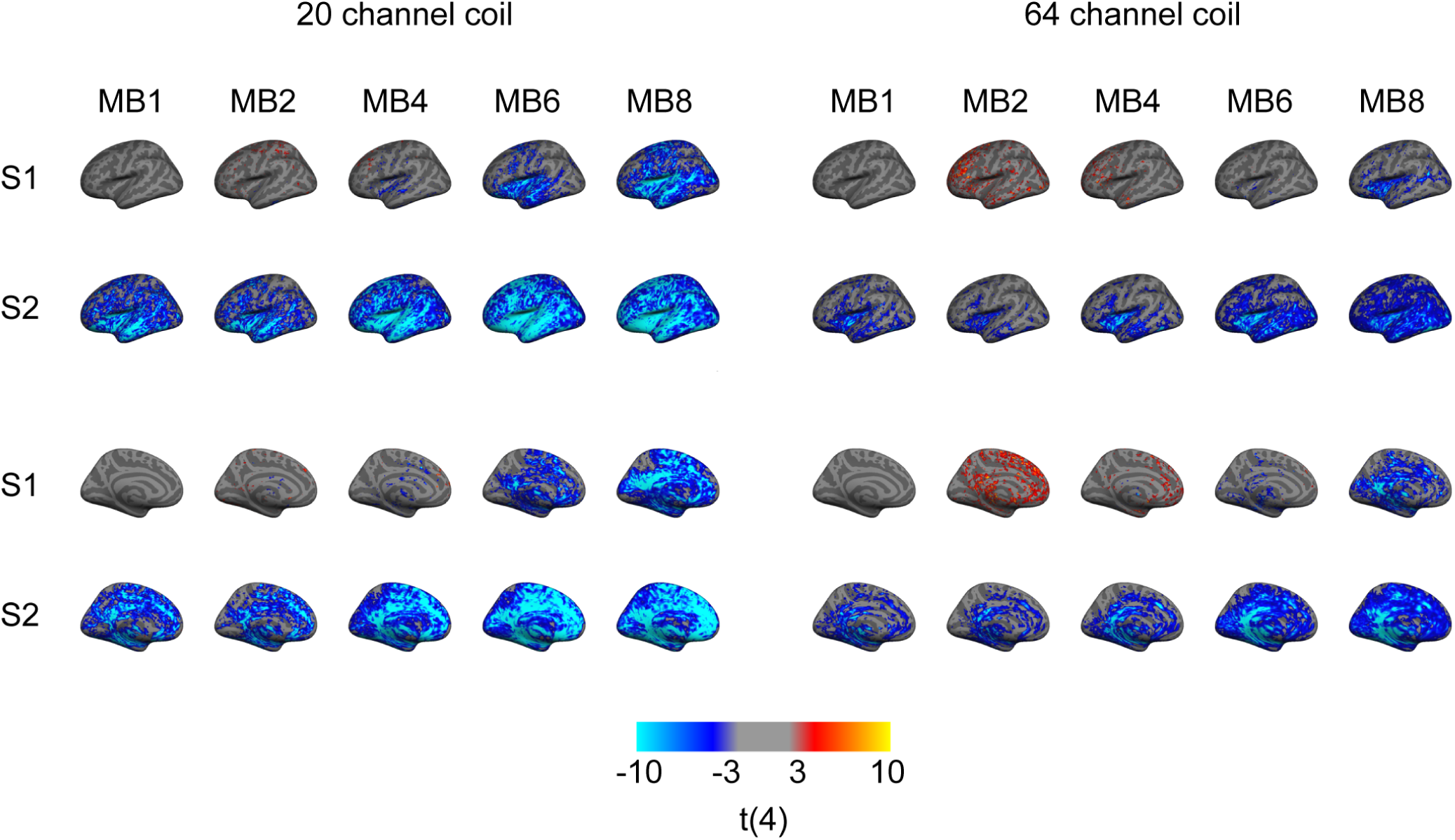
Uncorrected t-score maps (t_4_ = ±3, p < 0.02) of tSNR changes presented for each combination of acceleration factors compared to the baseline S1-MB1 (i.e., non-accelerated) projected onto a standard cortical surface for both the 20-channel coil (left) and 64-channel coil (right). Cold colors indicate a decrease in tSNR, while warm colors indicate an increase in tSNR. As predicted by the previous analyses, S-acceleration leads to the greatest—and most widespread—decreases in tSNR, though to a lesser extent when using the 64-channel coil. In all cases where tSNR does decrease, the temporal, insular, and medial regions appear most susceptible, in contrast to frontal and parietal regions.

Visualizing the results as the percentage of voxels with FWER-corrected tSNR decreases as a function of different acceleration methods partially reflects the results from the whole-brain average analysis. Namely, S-acceleration and higher acceleration factors, in general, tend to yield more voxels in which tSNR decreases. However, the voxelwise analysis may be more sensitive to the interaction between coil type and S-acceleration, as the 64-channel coil without S-acceleration tends to yield the least number of voxels with tSNR decreases (Fig. 3). This is especially evident at even the most liberal FWER-corrected threshold of p < ∼0.5 (Fig. 3, rightmost panel), in that combining the 64-channel coil with S1-MB8 produced a voxel count with tSNR loss that is less than 0.1% of the total voxel count, which is orders of magnitude smaller than other coil-acceleration combinations when comparing identical total acceleration. However, this acceleration combination was still outperformed by S1-MB4 and S1-MB6 (again, using the 64-channel coil), which produced the most favorable acceleration-to-tSNR-loss ratio and never surpassed a voxel count with tSNR loss of 0.001% (at any of the statistical thresholds).

### Temporal, medial, and insular regions suffer the greatest tSNR loss from acceleration

The uncorrected (from t_4_ = 3, p < 0.02) t-score maps echo the previous analyses (in that S-acceleration is more detrimental than MB-acceleration and that the 64-channel coil yields fewer tSNR losses) but additionally provide spatial information regarding trends in tSNR losses. In general, when we observed tSNR losses, the insular and temporal cortices along with medial regions tended to be most affected (Fig. 4). These losses were more pronounced when using the 20-channel coil and increased with increasing acceleration. On the other hand, the frontal and parietal cortices were less affected by tSNR losses (predominantly when using the 64-channel coil in combination with no S-acceleration and lower levels of MB-acceleration), and may have even yielded slight increases in tSNR.

## Discussion

This study investigated changes in tSNR resulting from applying acceleration methods to the acquisition of functional MRI data. Specifically, we aimed to probe which brain regions were most susceptible to tSNR loss stemming from slice-acceleration (specifically GRAPPA [7]), multiband-acceleration [14], and a combination thereof, when other scanning parameters were held constant, and whether the type of receiver coil additionally played a role underlying decreases in tSNR.

Overall, our findings were fourfold: 1) applying S-acceleration generally yielded lower levels of tSNR than applying MB-acceleration, 2) moderate levels of MB-acceleration without S-acceleration (e.g., S1-MB4 and S1-MB6) did not lead to a drastic loss of tSNR, 3) using the 64-channel coil tended to result in less tSNR decrease than using the 20-channel coil, and 4) the temporal, insular, and medial regions were most susceptible to decreases in tSNR when applying acceleration methods. Such results demonstrate the possibility that combining a more sensitive receiver coil with moderate MB-acceleration may lead to a favorable acceleration-to-tSNR-loss ratio in situations where accelerating a scan is necessary, depending on the brain regions under investigation.

These results are in line with previous research showing decreases in tSNR as a function of increasing acceleration factors [12, 18, 19], which we extended by additionally demonstrating differences between a 20-channel and a 64-channel coil via voxelwise analyses and, crucially, selecting scanning parameters that could function for all combinations of acceleration factors (with the exception of the number of slices for MB8); by otherwise maintaining constant parameters across conditions [20], the explanation of observed tSNR decreases is better constrained to changes in the acceleration factors themselves. Prior work has additionally demonstrated benefits in signal sensitivity when using MB-acceleration in certain scenarios [12, 18, 19, 22, 35], which, in our case, can also be seen as the modest tSNR increases in frontal regions at moderate MB-acceleration levels (Fig. 4). However, as caution has been advised in applying such acceleration methods [35], we would also take a more statistically conservative approach in seeking tSNR increases (or benefits in general) from pushing cutting-edge hardware techniques to their limits, as opposed to a more statistically liberal approach in detecting tSNR decreases (or detriments in general), which was the scope of the current work.

One issue that arises with respect to our statistical approach is that our sample size likely yielded overly conservative FWER corrected thresholds. With five subjects, the permutation space encompassed only 31 permutations, yielding more extreme minimum t-scores than a larger permutation space would. These extreme thresholds reduce the likelihood of detecting tSNR decreases, especially in moderate effect sizes; as such, the results presented in figure 3 should be seen as a lower-bound on the number of voxels expected to show decreases in tSNR.

Another more practical limitation in our approach is that, in order to use constant scanning parameters for all combinations of acceleration factors that we employed, it was necessary to use an unusually long TR of 4800 ms. For this reason, our findings may not quantitatively inform research that uses more “standard” TRs (e.g., in the range of 1000 – 2500 ms). However, using the long TR provided us with the benefit that, by being able to use otherwise constant scanning parameters, we could attribute changes in tSNR to changes in the acceleration factor. As such, the qualitative aspect of our results should nevertheless provide a reasonable guideline for researchers to consider when optimizing scanning protocols for their own scientific or clinical investigations, especially in the event that there are a priori brain regions to be studied. In fact, optimizing scanning protocols with acceleration methods has a clear benefit at the single-subject level. For example, tailoring a scanning protocol to an individual not only shifts the focus of the investigation to individual-specific information [36], but it can also reduce overall scan time (and thus the participant’s comfort) and potentially increase signal quality [18, 35]. These benefits are especially likely to express themselves during longitudinal studies.

In summary, we applied a combination of slice- and multiband-acceleration to functional MRI data acquisition (using otherwise constant sequence parameters) of both a phantom and human participants. Investigating the changes in temporal SNR as a function of such acceleration methods revealed a general decrease as the acceleration factor increases, but this decrease was more noticeable in S-acceleration compared to MB-acceleration. Moreover, moderate levels of MB-acceleration with no S-acceleration seemed not to lead to drastic losses of tSNR (i.e., S1-MB2 through S1-MB6), particularly when using a receiver coil with more elements (in our case, 64 channels), and especially in frontal and parietal regions of the brain. On the other hand, temporal, insular, and medial regions seemed to be most susceptible to decreases in tSNR. Our findings provide a guideline for researchers to consider when seeking to optimize their own research/clinical protocols depending on the experimental question and the available hardware.

## Author contributions

Experiment conception: JVS

Data acquisition: JVS, PS, and MT

Data analysis: JVS, PS, and SML

Initial manuscript draft: PS and SML

Manuscript revision and finalization: All authors

## Funding

MT is supported by a scholarship from the Universität Bayern e.V.

The formatting style for this manuscript is a derivative of the “HenriquesLab bioRxiv template” by Ricardo Henriques, used under CC BY 4.0.

## References

1. Mansfield, P. Multi-planar image formation using. J. Phys. C: Solid State Phys., 10:L55–L58, 1977.

2. Ogawa, S., Lee, T. M., Kay, A. R., and Tank, D. W. Brain magnetic resonance imaging with contrast dependent on blood oxygenation. Proceedings of the National Academy of Sciences, 87(24):9868–9872, dec 1990. doi: 10.1073/pnas.87.24.9868.

3. Belliveau, J., Kennedy, D., McKinstry, R., Buchbinder, B., Weisskoff, R., Cohen, M., Vevea, J., Brady, T., and Rosen, B. Functional mapping of the human visual cortex by magnetic resonance imaging. Science, 254(5032):716–719, nov 1991. doi1: 10.1126/science.1948051.

4. Kwong, K. K., Belliveau, J. W., Chesler, D. A., Goldberg, I. E., Weisskoff, R. M., Poncelet, B. P., Kennedy, D. N., Hoppel, B. E., Cohen, M. S., and Turner, R. Dynamic magnetic resonance imaging of human brain activity during primary sensory stimulation. Proceedings of the National Academy of Sciences of the United States of America, 89(12):5675–9, 1992.

5. Biswal, B., Zerrin Yetkin, F., Haughton, V. M., and Hyde, J. S. Functional connectivity in the motor cortex of resting human brain using echo-planar mri. Magnetic Resonance in Medicine, 34(4):537–541, oct 1995. doi: 10.1002/mrm.1910340409.

6. Feinberg, D. A., Hale, J. D., Watts, J. C., Kaufman, L., and Mark, A. Halving MR imaging time by conjugation: demonstration at 3.5 kG. Radiology, 161(2):527–531, nov 1986. doi: 10.1148/radiology.161.2.3763926.

7. Griswold, M. A., Jakob, P. M., Heidemann, R. M., Nittka, M., Jellus, V., Wang, J., Kiefer, B., and Haase, A. Generalized autocalibrating partially parallel acquisitions (GRAPPA). Magnetic resonance in medicine, 47(6):1202–10, 2002. doi: 10.1002/mrm.10171.

8. Pruessmann, K. P., Weiger, M., Scheidegger, M. B., and Boesiger, P. SENSE: Sensitivity encoding for fast MRI. Magnetic Resonance in Medicine, 42(5):952–962, nov 1999. doi: 10.1002/(SICI)1522-2594(199911)42:5<952::AID-MRM16>3.0.CO;2-S.

9. Sodickson, D. K. and Manning, W. J. Simultaneous acquisition of spatial harmonics (SMASH): Fast imaging with radiofrequency coil arrays. Magnetic Resonance in Medicine, 38(4):591–603, 1997. doi: 10.1002/mrm.1910380414.

10. Ohliger, M. A., Grant, A. K., and Sodickson, D. K. Ultimate Intrinsic Signal-to-Noise Ratio for Parallel MRI: Electromagnetic Field Considerations. Magnetic Resonance in Medicine, 2003. doi: 10.1002/mrm.10597.

11. Sodickson, D. K., Hardy, C. J., Zhu, Y., Giaquinto, R. O., Gross, P., Kenwood, G., Niendorf, T., Lejay, H., McKenzie, C. A., Ohliger, M. A., Grant, A. K., and Rofsky, N. M. Rapid Volumetric MRI Using Parallel Imaging With Order-of-Magnitude Accelerations and a 32-Element RF Coil Array. Academic Radiology, 12(5):626–635, may 2005. doi: 10.1016/j.acra.2005.01.012.

12. Chen, L., T. Vu, A., Xu, J., Moeller, S., Ugurbil, K., Yacoub, E., and Feinberg, D. Evaluation of highly accelerated simultaneous multi-slice EPI for fMRI. NeuroImage, 104:452–459, jan 2015. doi: 10.1016/j.neuroimage.2014.10.027.

13. Robson, P. M., Grant, A. K., Madhuranthakam, A. J., Lattanzi, R., Sodickson, D. K., and McKenzie, C. A. Comprehensive quantification of signal-to-noise ratio and g -factor for image-based and k -space-based parallel imaging reconstructions. Magnetic Resonance in Medicine, 60(4):895–907, oct 2008. doi: 10.1002/mrm.21728.

14. Nunes, R. G., Hajnal, J. V., Golay, X., and Larkman, D. J. Simultaneous slice excitation and reconstruction for single shot EPI. Proc. Intl. Soc. Mag. Reson. Med, 13(2):293, 2006.

15. Feinberg, D. A., Moeller, S., Smith, S. M., Auerbach, E., Ramanna, S., Glasser, M. F., Miller, K. L., Ugurbil, K., and Yacoub, E. Multiplexed Echo Planar Imaging for Sub-Second Whole Brain FMRI and Fast Diffusion Imaging. PLoS ONE, 5(12):e15710. dec 2010. doi: 10.1371/journal.pone.0015710.

16. Moeller, S., Yacoub, E., Olman, C. a., Auerbach, E., Strupp, J., Harel, N., and Uğurbil, K. Multiband multislice GE-EPI at 7 tesla, with 16-fold acceleration using partial parallel imaging with application to high spatial and temporal whole-brain fMRI. Magnetic Resonance in Medicine, 63(5):1144–1153, may 2010. doi: 10.1002/mrm.22361.

17. Uğurbil, K., Xu, J., Auerbach, E. J., Moeller, S., Vu, A. T., Duarte-Carvajalino, J. M., Lenglet, C., Wu, X., Schmitter, S., Van de Moortele, P. F., Strupp, J., Sapiro, G., De Martino, F., Wang, D., Harel, N., Garwood, M., Chen, L., Feinberg, D. A., Smith, S. M., Miller, K. L., Sotiropoulos, S. N., Jbabdi, S., Andersson, J. L., Behrens, T. E., Glasser, M. F., Van Essen, D. C., and Yacoub, E. Pushing spatial and temporal resolution for functional and diffusion MRI in the Human Connectome Project. NeuroImage, 80:80–104, oct 2013. doi: 10.1016/j.neuroimage.2013.05.012.

18. Todd, N., Josephs, O., Zeidman, P., Flandin, G., Moeller, S., and Weiskopf, N. Functional Sensitivity of 2D Simultaneous Multi-Slice Echo-Planar Imaging: Effects of Acceleration on g-factor and Physiological Noise. Frontiers in Neuroscience, 11, mar 2017. doi: 10.3389/fnins.2017.00158.

19. Todd, N., Moeller, S., Auerbach, E. J., Yacoub, E., Flandin, G., and Weiskopf, N. Evaluation of 2D multiband EPI imaging for high-resolution, whole-brain, task-based fMRI studies at 3T: Sensitivity and slice leakage artifacts. NeuroImage, 124:32–42, jan 2016. doi: 10.1016/j.neuroimage.2015.08.056.

20. Xu, J., Moeller, S., Auerbach, E. J., Strupp, J., Smith, S. M., Feinberg, D. A., Yacoub, E., and Uğurbil, K. Evaluation of slice accelerations using multiband echo planar imaging at 3T. NeuroImage, 83:991–1001, dec 2013. doi: 10.1016/j.neuroimage.2013.07.055.

21. Boyacioğlu, R., Schulz, J., Koopmans, P. J., Barth, M., and Norris, D. G. Improved sensitivity and specificity for resting state and task fMRI with multiband multi-echo EPI compared to multi-echo EPI at 7 T. NeuroImage, 119:352–361, oct 2015. doi: 10.1016/j.neuroimage.2015.06.089.

22. Preibisch, C., Castrillón G., J. G., Bührer, M., and Riedl, V. Evaluation of Multiband EPI Acquisitions for Resting State fMRI. PLOS ONE, 10(9):e0136961. sep 2015. doi: 10.1371/journal.pone.0136961.

23. Vu, A. T., Phillips, J. S., Kay, K., Phillips, M. E., Johnson, M. R., Shinkareva, S. V., Tubridy, S., Millin, R., Grossman, M., Gureckis, T., Bhattacharyya, R., and Yacoub, E. Using precise word timing information improves decoding accuracy in a multiband-accelerated multimodal reading experiment. Cognitive Neuropsychology, 33(3-4):265–275, may 2016. doi: 10.1080/02643294.2016.1195343.

24. Setsompop, K., Gagoski, B. A., Polimeni, J. R., Witzel, T., Wedeen, V. J., and Wald, L. L. Blipped-controlled aliasing in parallel imaging for simultaneous multislice echo planar imaging with reduced g-factor penalty. Magnetic Resonance in Medicine, 67(5):1210–1224, may 2012. doi: 10.1002/mrm.23097.

25. Smith, S. M., Jenkinson, M., Woolrich, M. W., Beckmann, C. F., Behrens, T. E. J., Johansen-Berg, H., Bannister, P. R., De Luca, M., Drobnjak, I., Flitney, D. E., Niazy, R. K., Saunders, J., Vickers, J., Zhang, Y., De Stefano, N., Brady, J. M., and Matthews, P. M. Advances in functional and structural MR image analysis and implementation as FSL. Neuroimage, 23 (SUPPL. 1):S208–S219, 2004. doi: 10.1016/j.neuroimage.2004.07.051.

26. Dale, A. M., Fischl, B., and Sereno, M. I. Cortical Surface-Based Analysis. NeuroImage, 9 (2):179–194, feb 1999. doi: 10.1006/nimg.1998.0395.

27. Fischl, B., Sereno, M. I., and Dale, A. M. Cortical Surface-Based Analysis. NeuroImage, 9 (2):195–207, feb 1999. doi: 10.1006/nimg.1998.0396.

28. Oosterhof, N. N., Connolly, A. C., and Haxby, J. V. CoSMoMVPA: multi-modal multivariate pattern analysis of neuroimaging data in Matlab / GNU Octave. Frontiers in Neuroinformatics, 10(July):047118, 2016. doi: 10.1101/047118.

29. Friedman, L. and Glover, G. H. Report on a multicenter fMRI quality assurance protocol. Journal of Magnetic Resonance Imaging, 23(6):827–839, jun 2006. doi: 10.1002/jmri.20583.

30. Smith, S. M. Fast robust automated brain extraction. Human Brain Mapping, 17(3):143–155, nov 2002. doi: 10.1002/hbm.10062.

31. Jenkinson, M. and Smith, S. A global optimisation method for robust affine registration of brain images. Medical Image Analysis, 5(2):143–156, jun 2001. doi: 10.1016/S1361-8415(01)00036-6.

32. Jenkinson, M. Improved Optimization for the Robust and Accurate Linear Registration and Motion Correction of Brain Images. NeuroImage, 17(2):825–841, oct 2002. doi: 10.1016/S1053-8119(02)91132-8.

33. Risk, B. B., Kociuba, M. C., and Rowe, D. B. Impacts of simultaneous multislice acquisition on sensitivity and specificity in fMRI. NeuroImage, 172:538–553, may 2018. doi: 10.1016/j.neuroimage.2018.01.078.

34. Nichols, T. E. and Holmes, A. P. Nonparametric permutation tests for functional neuroimaging: A primer with examples. Human Brain Mapping, 15(1):1–25, jan 2001. doi: 10.1002/hbm.1058.

35. Demetriou, L., Kowalczyk, O. S., Tyson, G., Bello, T., Newbould, R. D., and Wall, M. B. A comprehensive evaluation of increasing temporal resolution with multiband-accelerated protocols and effects on statistical outcome measures in fMRI. NeuroImage, 176(May): 404–416, aug 2018. doi: 10.1016/j.neuroimage.2018.05.011.

36. Dubois, J. and Adolphs, R. Building a Science of Individual Differences from fMRI. Trends in Cognitive Sciences, 20(6):425–443, jun 2016. doi: 10.1016/j.tics.2016.03.014.

